# Dominant and nondominant distal radius microstructure: predictors of asymmetry and effects of a unilateral mechanical loading intervention

**DOI:** 10.1101/2021.01.12.426389

**Authors:** Karen L. Troy, Megan E. Mancuso, Joshua E. Johnson, Tiffiny A. Butler, Bao Han Ngo, Thomas J. Schnitzer

## Abstract

Most information about distal radius microstructure is based on the non-dominant forearm, with little known about the factors that contribute to bilateral asymmetries in the general population, or what factors may influence bilateral changes over time. Here, we analyzed bilateral longitudinal high resolution peripheral quantitative computed tomography (HRpQCT) data collected over a 12-month period as part of a clinical trial that prescribed a well-controlled, compressive loading task to the nondominant forearm. Baseline data from 102 women age 21-40, and longitudinal data from 66 women who completed the 12-month trial, were examined to determine factors responsible for side-to-side asymmetries in bone structure and change in structure over time. Cross-sectionally, the dominant radius had 2.4%-2.7% larger cross-sectional area, trabecular area, and bone mineral content than the nondominant radius, but no other differences were noted. Those who more strongly favored their dominant arm had significantly more, thinner, closely spaced trabecular struts in their dominant versus nondominant radius. Individuals assigned to a loading intervention had significant bilateral gains in total bone mineral density (2.0% and 1.2% in the nondominant versus dominan sides), and unilateral gains in cortical area (3.1%), thickness (3.0%), bone mineral density (1.7%) and inner trabecular density (1.3%). Each of these gains were significantly predicted by loading dose, a metric that included bone strain, number of cycles, and strain rate. Within individuals, change was negatively associated with age, meaning that women closer to age 40 experienced less of a gain in bone versus those closer to age 21. We believe that dominant/nondominant asymmetries in bone structure reflect differences in habitual loads during growth and past ability to adapt, while response to loading reflects current individual physiologic capacity to adapt.

## 1. Introduction

Peak bone mass accrued during adolescence and young adulthood strongly influences bone health later in life. Lifetime peak bone mass is generally achieved by age 21 [1], with declines detectable after age 40[2]. Failure to reach optimal peak bone mass is considered a primary cause of osteoporosis. However, “optimal” is not well defined, and no target values exist. A previous theoretical study suggested that a 10% increase in peak bone mass would result in an additional 13 years of osteoporosis-free life [3]. We recently showed that a 1 SD increase in areal bone mineral density (aBMD) at the distal radius was associated with a 20% increase in bone stiffness [4]. Twenty to forty percent of peak bone mass is modifiable through lifestyle factors [2] such as sports participation and other bone-loading physical activity. These, combined with activities of daily living, influence distal radius density and strength. However, it is unclear how non-sports loading interacts with individual physiologic capacity to modify bone strength at this common fracture site. Studying asymmetries between dominant and nondominant arms may provide insights into this process.

Despite the growing body of literature on the importance of bone microstructure measures, only limited data have been reported in young healthy adult women[5], a key demographic for osteoporosis prevention, and the basis for calculating T-scores. Recent reports of racial and genetic differences in distal forearm microstructure [6] have highlighted the distinct mechanical contributions of the cortical versus trabecular compartments [7] to overall bone strength. These and other studies measure only the nondominant limb, based on previous DXA literature reporting side-to-side differences in forearm BMD as a result of habitual physical activity [8]. Most bilateral pQCT studies have focused on racquet [9] or throwing-sports [10] players, with an emphasis on documenting the effect of high intensity physical activity on side-to-side differences in bone structure. Despite these populations being interesting as models to understand bone adaptation, data collected in these athletes do not represent the general population. Additionally, hand dominance is a continuum and not a binary category, with over 35% of women being mixed-handed [11]. Because predominantly nondominant arms have been studied, little is known about how bone microstructure differs between dominant and nondominant arms in non-athlete populations, how these measures are related to habitual upper extremity use, are affected by handedness, or change over time. This is important because between-arm asymmetries reflect the structural adaptations that result from bone loading during asymmetrical activities of daily living. Furthermore, these structural differences may be an indicator of an individual’s physiologic capacity for adaptation. For example, individuals with greater physiologic capacity to adapt could express larger asymmetries for a given set of asymmetrical bone loading activities. On the other hand, a unilateral loading activity might induce changes to the non-loaded arm due to systemic factors.

In a clinical trial that used a voluntary compressive distal radius loading activity, we recently reported that 10-15% of the 12-month changes in the nondominant (loaded) distal radius BMC could be explained by loading dose, a linear combination of the peak strain, strain rate, and number of loading cycles [12]. Here, we analyzed longitudinal high resolution peripheral quantitative computed tomography (HRpQCT) data collected over a 12-month period as part of this parent study. The present analysis sought to answer the following questions, in a group of healthy women age 21-40: (1) What is the relationship between handedness, physical activity history, and bilateral distal radius microstructure? (2) What factors influence individual change in dominant and nondominant distal radius microstructure during the intervention (month 0-12)? We hypothesized that individuals who were more mixed-handed, participated in both-handed physical activities, and had similar grip strength on each side would have fewer side-to-side differences in bone microstructure than those with large asymmetries in hand use or strength. We expected that age, blood levels of 25,OH vitamin D, and baseline values of bone mass would predict bilateral changes to the distal radius, with loading dose affecting only the nondominant (loaded) side during the observation period.

## 2. Methods

Healthy women aged 21-40 were recruited to participate in a prospective clinical trial investigating bone adaptation and mechanical strain environment (NCT04135196). Subject recruitment, inclusion criteria, and screening are described in detail elsewhere [12]. Briefly, women responding to online advertisements were contacted and screened via telephone survey. Individuals were excluded if they had irregular menstrual cycles, body mass indices outside the range 18–25□kg/m^2^, no regular calcium intake, or reported taking medications known to affect bone metabolism. Those regularly participating (> 2 time per month) in sports that apply high-impact loads to the forearm (e.g. gymnastics, volleyball) were also excluded. Qualified subjects had 25-hydroxyvitamin D serum above 20□ng/ml and a total forearm DXA T-score between −2.5 and 1.0. Data for qualified subjects were collected either during the screening or a single visit within approximately two weeks of screening. All participants provided written, informed consent to the institutionally approved study between January 2014 and June 2017.

The loading intervention is described in detail elsewhere [12], and consisted of voluntary compressive loading of the nondominant forearm arm for 100 cycles/day, 3 days/week, for 12 months. Participants were prospectively assigned to one of four loading groups that varied the magnitude and rate of the force, or were assigned to a non-intervention control group. In total, 102 women age 21-40 were enrolled and completed baseline testing and 66 completed the 12-month intervention. Of these individuals, 13 were assigned to a non-loading control group, while the remaining 53 were assigned a loading intervention.

### 2.1 Demographics and Bone Loading

As previously described [12], height, weight, and grip strength were measured using a wall-mounted stadiometer, an analog scale, and a hydraulic hand-grip dynamometer (Baseline; White Plains, NJ). Average daily calcium intake (mg/day) was estimated using a 10-item questionnaire that tallied weekly consumption of calcium-containing foods and beverages [13].

Forearm mechanical loading was measured in three ways to reflect activities of daily living (handedness), physical activity, and the intervention. Handedness was determined using the Edinburgh Index [14], a validated questionnaire that assigns a laterality index score ranging from −100 (completely left-hand dominant), 0 (completely mixed-handed) to +100 (completely right-hand dominant). Forearm loading due to physical activity was estimated using a site-specific arm bone loading index (*armBLI*) algorithm [15], based on activity histories collected using a validated survey[16]. For the purposes of calculating *armBLI* scores, individuals were considered either right or left-handed in a binary fashion. The *armBLI* algorithm scores activities based on the magnitude, rate, and frequency (days/week) of loads applied to the nondominant arm as:

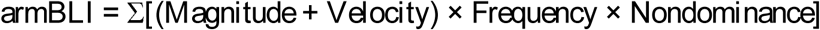

where the nondominance multiplier corrects for activities preferentially loading the dominant arm. The multiplier is 0.33 for predominantly unilateral activities (e.g., tennis), 0.66 for somewhat unilateral activities (e.g. softball), and 1.0 for bilateral activities (e.g. gymnastics). ArmBLI was similarly calculated for the dominant arm, but with the non-dominance multiplier set to 1.0. Here we report the total average annual armBLI score (armBLI/year). Loading dose from the intervention was determined based on force parameters from load cell recordings that were converted to bone strain using validated [17] subject-specific finite element models. Loading dose was calculated as: mean(Peak-to-peak strain magnitude)*mean(Strain rate)*[#bouts], and was expressed in units of με^2^*s^-1^*bouts*10^−5^.

### 2.2 High Resolution pQCT

High-resolution peripheral quantitative computed tomography (HRpQCT; XtremeCT, Scanco Medical; Brüttisellen, Switzerland) scans of the distal radius in both the dominant and nondominant arms were performed according to the manufacturer’s standard *in vivo* scanning protocol. The scans consisted of 110 slices with an isotropic voxel size of 82□μm, encompassing a 9.02□mm axial region beginning 9.5□mm proximal to a reference line placed at the distal endplate. All scans were performed by trained technicians, and daily and weekly quality control scans were performed. Each scan was graded for motion on a scale from 1 (no motion) to 5 (severe motion artifact) [18], and only scans scoring 3 or better were included in the analysis.

HRpQCT scans acquired at 12-months were registered to the baseline scan using the manufacturer’s 2D region-matching algorithm. Follow-up scans were only included if they overlapped at least 70% with the baseline region. Total mean cross-sectional area (CSA; mm^2^) and total volumetric bone mineral density (Tt.BMD; mgHA/cm^3^) were measured. Baseline total bone mineral content (Tt.BMC; g) was calculated as the product of these two metrics and the scan length at baseline (9.02 cm). Both cortical and trabecular cross-sectional area (Ct.A and TbA; mm^2^) were also calculated. Trabecular number (Tb.N; mm^−1^), thickness (Tb.Th; mm), spacing (Tb.Sp; mm), bone volume fraction (Tb.BVTV; cm^3^/cm^3^) and BMD (Tb.BMD; mgHA/cm^3^) were measured using the manufacturer’s standard analysis protocol. The trabecular region was further divided into inner (central 60%; Tb.BMDinn) and outer regions (outer 40%; Tb.BMDmeta). In our lab, the coefficient of variation (CV) for densitometric variables is < 0.3%. Cortical vBMD (Ct.BMD; mgHA/cm^3^) and cortical thickness (Ct.Th; mm) were calculated using the dual-threshold method [19, 20]. The CVs of these variables range from 0.4-4%. Parameters and 12-month changes were calculated based on the overlapping region.

### 2.3 Statistical Analysis

The normality of each measured variable was assessed by visual inspection of histogram distributions. Dominant versus nondominant bone structure were initially compared with paired t-tests. To assess demographic predictors of baseline bone structure, correlations were calculated between demographic and loading variables versus baseline HRpQCT values. To determine whether loading asymmetry is associated with greater between-arm bone differences at baseline, correlations were also calculated between percent differences in HRpQCT values (100%*(dominant – nondominant)/(mean of both sides)) versus percent differences in grip strength, armBLI scores, and versus the absolute value of laterality index.

To identify which HRpQCT variables changed over the 12-month observation period, we performed paired t-tests to compare baseline and 12-month values for each microstructural variable in each arm. Because we previously observed increases in some experimental groups [12], we first separated participants into two broad groups for the paired t-tests: those who were assigned to any mechanical loading intervention (n=53), and those that were not (n=13). To statistically control for the large number of comparisons being made this way, we adjusted our significance criterion using Holm’s Step-down Method [21]. Next, to identify predictors of 12-month change within the entire cohort, we examined the degree to which time (baseline, 12 months) and loading dose (a continuous variable, treated as a covariate) were related each bone parameter using a mixed linear model with a random intercept term for each participant and time being a repeated measure. Each hand was examined separately, with dominant hands and control subjects being assigned a loading dose of zero. For variables shown to change over time (significant effect of time or time*dose), additional regression models were fit to identify which demographic or loading variables may influence these changes. First, an initial model was generated with the dependent variable of 12-month change and predictors of baseline value and loading dose (based on our previous findings [12]). Then, a full model was generated that included all of the candidate variables (baseline value, loading dose, age, height, weight, vitamin D, calcium intake, armBLI, and grip strength). Lastly, a final model was generated after eliminating all non-significant predictors from the full model. An alpha level of 0.05 was used to detect significance. All statistical analyses were performed using SPSS v22.0.

## 3. Results

Most variables were normally distributed, although dominant/nondominant differences were not. Accordingly, side-to-side differences were assessed using Spearman (non-parametric) correlations, rather than Pearson correlations. Demographics and baseline microstructural variables are summarized in Table 1. In general, dominant and nondominant bone parameters were not consistently different, except for increased CSA, trabecular area, and total BMC in the dominant side, indicative of an overall larger bone morphology.

**Table 1a:** Demographics of the study participants

| Subject Characteristics | Mean | SD |
| --- | --- | --- |
| Age (years) | 28.4 | (5.6) |
| Height (cm) | 164.8 | (6.4) |
| Body Mass (kg) | 64.3 | (8.7) |
| Serum Vitamin D (ng/mL) | 31.7 | (9.5) |
| Calcium intake (mg/day) | 676.5 | (424.3) |
| Laterality index (Absolute decile) | 79 | (22) |
| Right/Left handers | 91/11 |  |

**Table 1b:** Baseline microstructural variables in the dominant and nondominant hands. The reported P-values are from paired t-tests.

|  | Non-Dominant |  | Dominant |  | Within-subject % difference |  | P-value |
| --- | --- | --- | --- | --- | --- | --- | --- |
|  | Mean | SD | Mean | SD | Mean | SD |  |
| **Grip strength** | 259.01 | 51.85 | 275.21 | 54.3 | 6.1 | 13.6 | **<0.001** |
| **armBLI** | 976.2 | 819.8 | 1117.7 | 946.2 | 12.6 | 15.0 | **<0.001** |
| **mean CSA (mm2)** | 270.08 | 43.92 | 276.37 | 42.3 | 2.4 | 8.2 | **0.008** |
| **Tt.BMD (mg/cm3)** | 303.41 | 50.37 | 301.82 | 46.90 | -0.4 | 8.0 | 0.540 |
| **Tt.BMC (g)** | 72.62 | 11.14 | 74.17 | 11.79 | 2.0 | 5.5 | **<0.001** |
| **Ct.Area (mm2)** | 47.17 | 9.33 | 47.73 | 9.49 | 1.1 | 11.8 | 0.311 |
| **Tb.Area (mm2)** | 217.52 | 44.56 | 222.72 | 42.47 | 2.7 | 10.9 | **0.032** |
| **Ct.BMD (mg/cm3)** | 864.42 | 54.68 | 861.78 | 50.51 | -0.3 | 3.9 | 0.438 |
| **Ct.Th (mm)** | 0.682 | 0.145 | 0.681 | 0.141 | -0.3 | 14.3 | 0.888 |
| **Tb.BMD (mg/cm3)** | 162.47 | 30.93 | 163.60 | 31.07 | 0.7 | 6.6 | 0.438 |
| **Tb.BMDmeta (mg/cm3)** | 221.54 | 30.10 | 222.47 | 30.63 | 0.4 | 4.3 | 0.338 |
| **Tb.BMDinn (mg/cm3)** | 121.67 | 32.63 | 122.94 | 32.66 | 1.1 | 11.7 | 0.296 |
| **Tb.BVTV (cm3/cm3)** | 0.135 | 0.026 | 0.136 | 0.026 | 0.7 | 6.5 | 0.272 |
| **Tb.N (mm-1)** | 2.024 | 0.254 | 2.038 | 0.249 | 0.7 | 8.6 | 0.433 |
| **Tb.Th (mm)** | 0.067 | 0.010 | 0.067 | 0.010 | -0.1 | 9.5 | 0.816 |
| **Tb.Sp (mm)** | 0.434 | 0.068 | 0.432 | 0.067 | -0.8 | 8.9 | 0.539 |

### 3.1 Predictors of Baseline Bone Structure

In both arms, bone cross sectional area and trabecular area were positively correlated with grip strength and body size (height and weight). Increasing age was associated with increases in trabecular spacing and decreases in trabecular number. We also observed a negative relationship between cortical density and grip strength (Table 2). We did not observe any relationship between bone microstructure and vitamin D status, calcium intake, or armBLI in either arm.

**Table 2:** Pearson correlations between demographic factors and bone structure. Significant values are shown in bold font and indicated with an asterisk.

|  | CSA |  | Tt.BMC |  | Ct.BMD |  | Tb.Area |  | Tb.N |  | Tb.Sp |  |
| --- | --- | --- | --- | --- | --- | --- | --- | --- | --- | --- | --- | --- |
|  | ND | D | ND | D | ND | D | ND | D | ND | D | ND | D |
| Age | -0.038 | 0.064 | 0.044 | 0.023 | 0.178 | 0.038 | -0.068 | 0.047 | **-0.235\*** | -0.195 | **0.207\*** | **0.209\*** |
| Grip strength | **0.530\*\*** | **0.495\*\*** | **0.354\*\*** | **0.224\*** | **-0.268\*\*** | **-0.211\*** | **0.501\*\*** | **0.426\*\*** | 0.104 | -0.026 | -0.027 | 0.026 |
| Height | **0.536\*\*** | **0.545\*\*** | **0.281\*\*** | **0.205\*** | **-0.231\*** | **-0.349\*\*** | **0.509\*\*** | **0.534\*\*** | 0.051 | 0.100 | -0.020 | -0.037 |
| Weight | **0.352\*\*** | **0.409\*\*** | **0.338\*\*** | **0.346\*\*** | 0.034 | -0.043 | **0.310\*\*** | **0.362\*\*** | 0.187 | **0.251\*** | -0.171 | **-0.212\*** |

\*p<0.05 \*\*p<0.01

### 3.2 Predictors of Side-to-Side Differences in Bone Structure

Laterality index was weakly associated with differences in trabecular microstructure. More use/reliance on the dominant hand was associated with relatively larger dominant vs. nondominant Tb.N (r=0.252 p=0.029), smaller Tb.Th (r=-0.224, p=0.029) and smaller Tb.Sp (r=− 0.230, p=0.025; Figure 1). That is, those who more strongly favor their dominant arm tended to have more, thinner, closely spaced trabecular struts in their dominant versus nondominant radius. Asymmetries in grip strength and armBLI were not related to microstructural differences.

**Figure 1:**
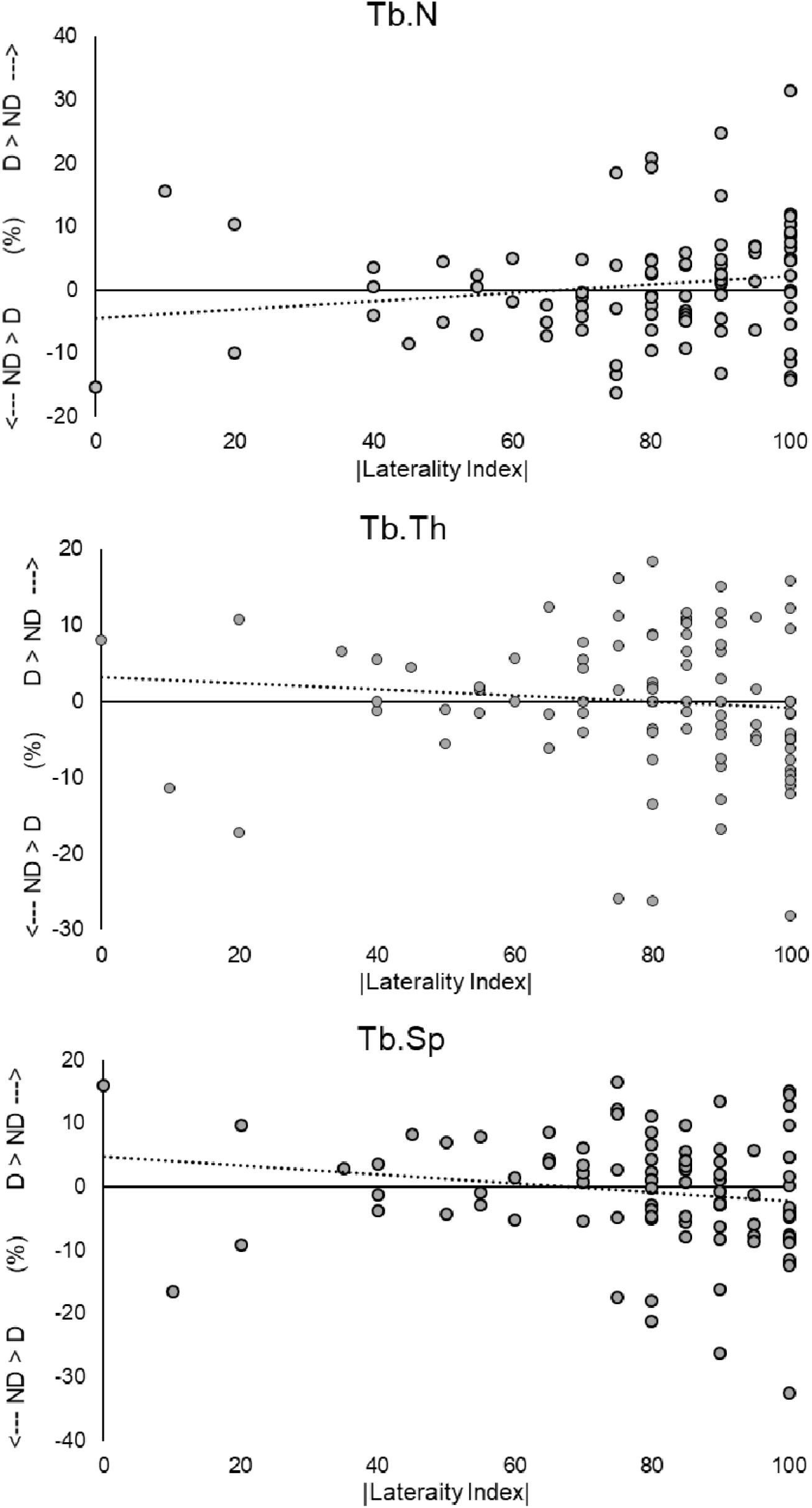
Dominant - Nondominant differences (expressed as a percentage) of (a) Trabecular Number, (b) Trabecular thickness, and (c) Trabecular spacing versus the absolute value of laterality index. Higher values of laterality index indicate high reliance on the dominant hand. This measure of laterality index does not depend on handedness. Dashed lines show the best-fit line, which was significant in each of these plots.

### 3.3 Predictors of 12-Month Change in Dominant and Nondominant Radii

Twelve-month changes to bone were greatest in the nondominant (loaded) side of individuals who were assigned an experimental loading intervention. In this group, the nondominant side bone experienced structural improvements that included increases to Tt.BMD, Tb.BMDinn, Ct.A and Ct.BMD (Figure 2a). In the dominant (within-subject control) arm, only Tt.BMD increased, suggesting that most benefits of the intervention are limited to the loaded side only. Individuals assigned to the control group experienced few significant changes (only increases to Ct.BMD; Figure 2b).

**Figure 2:**
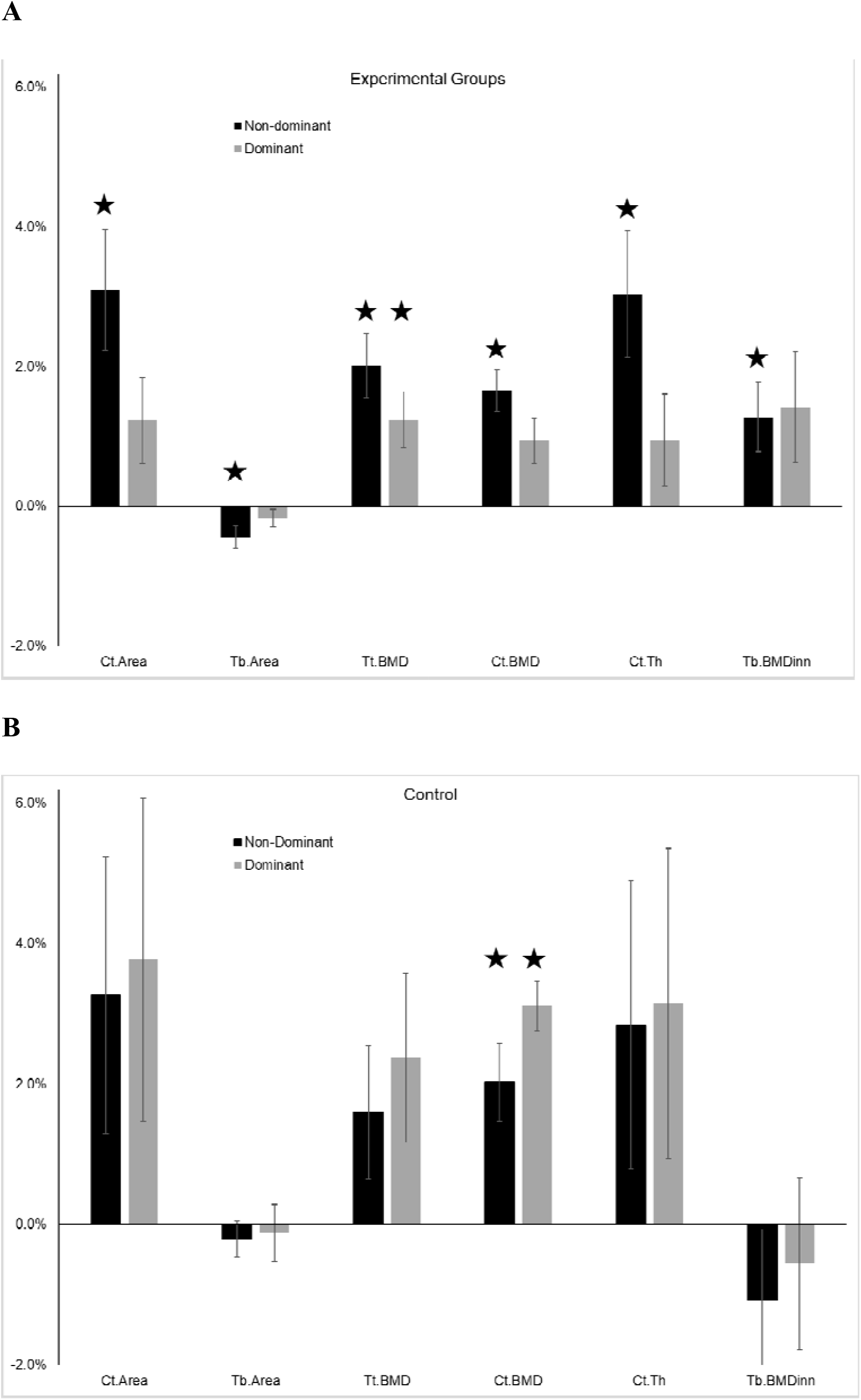
Dominant and nondominant changes in distal radius microstructure over a 12-month period. (A) shows all individuals who were assigned to a unilateral loading intervention on the nondominant side (n=53). (B) shows individuals who were assigned to a non-loading control group (n=13). Variables that changed significantly over the observation period are indicated with a★. Error bars show standard error.

With all participants combined, we observed significant time-related increases in Tt.BMD, Ct.A, and Ct.BMD in the nondominant side (Table 3). We also observed positive significant time*dose interactions in Tt.BMD, Tb.BMD, Tb.BMDinn, and Tb.BVTV. Tb.A was positively associated with dose, but we detected no time or time*dose changes for this variable. No significant time, dose, or time*dose relationships were observed for any variables in the dominant hand. This is consistent with the finding that few changes were observed in the dominant arm and the definition of dose as zero for the dominant arm.

**Table 3:** (top) Regression models to examine time and dose-related changes to microstructural and density variables. Only the nondominant side is shown here, because no models were significant for the dominant side. (bottom) Regression models to determine factors that predict 12-month change in each variable. In all models, significant coefficients and models are shown in bold.

| | Dependent Variable | $\beta_1$ (time) | $\beta_2$ (dose) | $\beta_3$ (time*dose) |
| --- | --- | --- | --- | --- |
| [Var]=$\beta_1$*time + $\beta_2$*dose+ $\beta_3$*time*dose+$\epsilon_i$ (ND side) | Tt.BMD | **3.710** | 1.05E-02 | **4.29E-03** |
|  | Ct.A | **1.010** | -1.65E-03 | 5.87E-04 |
|  | Tb.A | -0.067 | **1.92E-02** | 7.14E-04 |
|  | Ct.BMD | **12.180** | 1.16E-03 | 4.12E-03 |
|  | Tb.BMD | 0.004 | -4.57E-03 | **2.06E-03** |
|  | Tb.BMDinn | 0.050 | 4.26E-03 | **2.61E-03** |
|  | Tb.BVTV | 0.000 | -3.73E-06 | **1.87E-06** |

|  | Dependent Variable | Adj. R<sup>2</sup> | Final Regression Equation |
| --- | --- | --- | --- |
| Predictors of 12-month change (ND side) | ΔTt.BMD | **0.106** | **-0.299(age)+5.04E-3(dose)+0.001(armBLI)+3.756** |
| | ΔCt.A | **0.041** | **-0.109(age)-0.001(calcium)+0.001(armBLI)+4.548** |
| | ΔCt.BMD | **0.189** | **-0.197(baseline)+6.63E-3(dose)+103.5** |
| | ΔTb.BMD | **0.061** | **2.04E-3(dose)+0.002** |
| | ΔTb.BMDinn | **0.198** | **0.039(baseline)+3.22E-3(dose)-4.62** |
| | ΔTb.BVTV | **0.063** | **1.73E-6(dose)-0.00006** |

Bold For R<sup>2</sup> indicates p<0.05 for F-test of overall model fit
Bold For $\beta_1$ or $\beta_2$ indicates p<0.05 for t-test of significance for coefficient in the final model

Based on these findings, our regression analyses to determine predictors of 12-month change were limited to the nondominant side. On this side, dose was a positive significant predictor of 12-month change for all of the variables tested except for Ct.A. Baseline value was negatively associated with change in Ct.A, but positively associated with change in Tb.BMDinn. In addition to dose and baseline value, final models for Tt.BMD and Ct.A included negative coefficients for age and positive coefficients for armBLI, although the t-statistics for these coefficients did not reach significance in the final models (p=0.06 to 0.09; Table 3). Overall, these regression models were able to explain 4% to 20% of the total variance in the outcome variables of interest.

## 4. Discussion

Our purpose was to determine factors that could explain between- and within-subject variation in bone structure, and to identify predictors of change over a 12-month observation period. We analyzed baseline and 12-month changes in distal radius microstructure in the dominant and nondominant hands of 21-40 year old women. Between individuals, we found that grip strength and body size were positively associated with multiple measures of dominant and nondominant bone density and structure, and that trabecular microstructure was negatively correlated with age, even within our age range of 21-40 years. Upper extremity bone loading associated with physical activity (armBLI), calcium intake, and serum levels of 25-OH Vitamin D were not related to any measures of bone structure or density. Within individuals, we found that dominant/nondominant differences weakly depended on laterality index, especially in the trabecular bone compartment. Finally, we found that significant 12-month changes occurred in the nondominant arm, but not the dominant arm, of those assigned a unilateral loading intervention. Change in the nondominant arm was positively related to the loading dose provided by the intervention. In addition, some change variables were related to baseline values, armBLI, age, and calcium intake.

Our findings that grip strength and body size were most strongly related to distal radius structure are consistent with previous reports, which have documented strong relationships between strength, body size, and distal radius BMD (e.g. [22]). We previously reported that nondominant distal radius cross-sectional area was positively correlated with age, height, and grip strength in a large subset of this cohort [4]. Here, we confirm that these relationships are similar in both the dominant and nondominant forearms. We observed negative relationships between Ct.BMD, grip strength, and height. This is also consistent with our previous reports of grip strength and height being positively related to cortical porosity [4]. Because our HRpQCT scan region of interest was located at a fixed distance from the distal subchondral plate of the radius, scans from taller individuals are located proportionally more distally, where porosity is higher. The association between scan location, cross-sectional area, and cortical density has been previously reported [23], though it is unclear how grip strength is involved. Also similar to our previous report of diminished Tb.BMD with age, we observed that among adults (age 21-40) who are at peak bone mass, there was a negative correlation with age (between subjects) but no within-subject decreases over 12 months in trabecular bone.

The differences we observed in dominant versus nondominant forearms are smaller than reports in sedentary women based on DXA data[8], which reported 4% higher BMD and 10% higher BMC in the dominant forearm. Our data show that dominant and nondominant distal radii differ primarily in cross-sectional area, with nondominant radii having 2.7% smaller Tb.A compared to dominant radii. Consistent with our hypotheses, we found that heavy reliance on the dominant hand (with large absolute values for Laterality Index) was weakly associated with increased asymmetries in trabecular microstructure. The Edinburgh Index is a well-validated tool for quantifying handedness, but does not explicitly provide information about how this affects upper extremity bone loading. Individuals often choose their dominant hand for dexterity tasks such as manipulating an object, but their nondominant hand for stabilization tasks such as holding the object, highlighting that even highly lateralized individuals likely load both hands regularly. Sports, on the other hand, are frequently more unilateral. As such, others have examined side-to-side differences in athletic populations that preferentially load the dominant arm, such as baseball and racquet sports. In these athletes, differences in regional BMC as large as 28% have been documented [10]. Our study excluded individuals who participated regularly in sports with heavy upper extremity loading, so armBLI in the present study may underestimate the magnitude of these sports-driven effects. Because physical activity early in life tends to increase bone size[24], while activity later in life has a greater effect on density and cortical thickness[9, 12], the side-to-side differences that we observed here may be an indicator of asymmetries in habitual loads that are not captured in the armBLI or Laterality Index measures. For an idealized circular cross-section, a 2.7% increase in cross-sectional area represents an 11% increase in resistance to bending, highlighting the mechanical importance of these small differences.

We observed significant increases to inner trabecular bone density and cortical bone density and quantity in the nondominant side of individuals who participated in the unilateral loading intervention. We also observed significant bilateral increases to Tt.BMD in the loading groups. In general, changes in the nondominant (loaded) radius variables were about twice as large as the dominant (within-subject control) radius. It is likely that the changes we observed reflect a combination of both local and systemic mechanisms. The more robust changes in the non-dominant arm are likely related to local bone loading. We previously showed that a similar mechanical loading regimen elicited changes in distal radius BMD that were related to local mechanical strain environment [25]. In a subset of this cohort, we showed that local trabecular strain magnitude and gradient were higher near formation versus resorption sites [26]. Similarly, local trabecular strain environment has been linked to local formation/resorption in small animals [27]. Mechanical loading also activates systemic biochemical signals, including short-term increases in circulating parathyroid hormone [28]. This systemic response increases bone turnover, resulting in short-term gains to bone mass throughout the body [29]. Bilateral gains in total bone density may also be related to neuronal activity, which has been shown to influence bone adaptation [30], and elicit bilateral muscle gains following unilateral strength training interventions [31]. Unilateral *in vivo* small animal loading models routinely report increases in the unloaded limb, although many of these animals continue to grow during the intervention period (e.g. [32]).

Our hypothesis that calcium intake and vitamin D levels would be related to baseline and change in bone was not generally supported. We excluded participants who had blood levels of 25-OH Vitamin D less than 20 ng/ml, so lack of range in this variable may explain the negative finding. When we previously examined factors associated with nondominant bone structural behavior in a large subset of this dataset [4] we also noted a lack of association with vitamin D. We similarly excluded individuals who reported no or very little dietary calcium intake (e.g. avoidance of dairy products). We observed that calcium intake was positively associated with nondominant change in Ct.A but not CSA, suggesting greater dietary calcium is associated with endosteal bone apposition following loading. This is consistent with a study showing that in calcium deficient mice, calcium repletion was associated with increased endosteal mineral apposition, but had no effect on the periosteal surface [33].

Our study had several important limitations. We studied a group of healthy adult women with no specialized sports history, dietary restrictions, or other qualities. Our cohort was representative of the population in a predominantly white, North American setting. Because factors such as genetics, diet, and lifestyle affect bone, our findings may not apply to other groups. The longitudinal study had a relatively high drop-out rate, which could bias some of the results. However, the characteristics of the participants who completed the study were not different from those who dropped out[12]. The study also had several important strengths. We used a comprehensive set of measures to characterize handedness and habitual upper extremity bone loading. We also report longitudinal changes to bone structure during a well controlled intervention, using both within- and between-subjects controls.

In summary, we analyzed longitudinal HRpQCT data from bilateral distal radii in healthy women age 21-40 to determine relationships between handedness, physical activity history, and bilateral asymmetries in bone structure, as well as baseline and longitudinal change in bone structure. We found that cross-sectional area was 2.4% larger in the dominant versus nondominant side, with corresponding differences in trabecular area and total BMC. Consistent with previous reports, we observed positive relationships between grip strength, height, weight, and bone size. We used continuous measures of handedness and upper extremity bone loading physical activity to examine, for the first time, how asymmetries in these parameters affected asymmetries in distal radius bone structure. We found that the degree of hand dominance was weakly related to asymmetries in trabecular microstructural parameters. We believe that dominant/nondominant asymmetries in bone structure reflect differences in habitual loads during growth and historical capacity to adapt, while bilateral response to the loading intervention is an indicator of current individual physiologic capacity to adapt. However, none of the metrics that we examined were strong predictors of these asymmetries. Finally, we observed that a 12-month unilateral distal radius loading intervention was associated with significant bilateral increases to total BMD, but that the effect in other variables was unilateral and consistently scaled with loading dose. Despite observing cross-sectional age-related declines to trabecular microstructure in the cohort at baseline, the longitudinal data showed little change in bone parameters in this group of women at peak bone mass.

## Acknowledgements

This research was fully supported by NIAMS of the National Institutes of Health under award number R01AR063691.The content is solely the responsibility of the authors and does not necessarily represent the official views of the National Institutes of Health. This material is also based upon work supported by the National Science Foundation Graduate Research Fellowship Program under Grant No. DGE□1106756. We thank Sabahat Ahmed for her organization and dedication as research coordinator, and Julie Teavanen, Nour Krayem, and Nicole Zaino for assistance in analyzing HRpQCT data.

Authors’ roles: Study conceived by KLT and designed by KLT with assistance from TJS. Data collection by MEM, KLT, JEJ, and TAB. Data analysis and interpretation: MEM, KLT, JEJ, BHN, TAB, TJS. Manuscript writing: KLT and MEM. Manuscript approval: MEM, KLT, JEJ, TAB, TJS.

